# Fermentation and its effect on the physicochemical and sensory attributes of cocoa beans in the Colombian Amazon

**DOI:** 10.1101/2024.06.24.600490

**Authors:** Andrés Felipe Ramírez González, Gustavo Adolfo Gutiérrez García, Paola Andrea Polanía-Hincapié, Luis Javier López, Juan Carlos Suárez

## Abstract

Cocoa (*Theobroma cacao* L.) is the basic raw material to produce chocolate and other derivatives such as cocoa butter, cocoa powder and cocoa liquor (cocoa paste), which requires a fermentation process that affects its chemical composition and sensory profile. The objective of this study was to monitor the biochemical, physical and sensory changes during fermentation of cocoa beans in cocoa bean processing plants in the department of Caquetá, Colombia. During fermentation, the temperature of the mass and the pH of the pulp and beans were monitored at the different cocoa processing plants (sites ASOACASAN ASA, COMICACAO CMI, COMCAP COC). Also, at two points during fermentation (days 4 and 7), physical properties of the bean were determined, such as variables related to bromatological composition, polyphenolic compounds and antioxidant activity as sensory attributes at the different sites. An increase in dough temperature was found, however the pH of the cotyledon decreased during the fermentation process and the fat and moisture content varied with fermentation time. At the site level, total polyphenol content (TPC), total flavonoids (TF), DPPH and FRAP contents were statistically different, with COC being different from the other sites. The TPC was higher at the COC site (507 mg GAE/g Cocoa) with respect to the other sites (< 360 mg GAE/g Cocoa). The TF content followed a similar behavior to TPC, with significant differences between sites and differences between fermentation times for ASA. The TF was higher in COC (309.1 mg Catechin/g cocoa) with respect to CMI (215.6 mg Catechin/g cocoa) and ASA (185.7 mg Catechin/g cocoa). Values in DPPH ranged from 5869.3 to 7781.8 𝜇mol Trolox/g cocoa and for the FRAP assay ranged from 369.8 to 606.7 mg AA/g cocoa among the sites.

## 1. Introduction

Cocoa (*Theobroma Cacao* L.) is the basic raw material to produce chocolate and other derivatives such as cocoa butter, cocoa powder and cocoa liquor (cocoa paste) [1], which has sustained part of the daily diet of people and is currently making inroads in the cosmetics, perfumery and pharmaceutical industries [2] with a requirement in the final quality of the product. Recent studies have reported that grain quality in sensory terms is affected by different factors such as the genetics of the material [3], environmental conditions [4], the state of maturity of the grain and practices in the postharvest stage [5], such as those related to fermentation and drying [6,7].

The composition and quality of cocoa beans is determined by a set of physical, chemical and sensory qualities [8]. According to García et al. [9] the main biochemical changes that influence the quality of cocoa beans occur during the fermentation stage, since it is there where different microbial communities (yeasts, lactic acid and acetic acid bacteria) intervene in the metabolization of sugars and other compounds [10]. Fermentation is a process in which fresh cocoa beans are placed in a wooden box for four to seven days. During this process, the microorganisms metabolize the pulp that surrounds the beans and generate reactions that result in an increase in temperature, a decrease in pH and death of the embryo [11]. During fermentation, different biochemical reactions occur on various substances, including antioxidants, polyphenols and methylxanthines [1]. For example, polyphenols are products of the secondary metabolism of plants, characterized by aromatic rings with hydroxyl groups as substituents, which are directly related to the sensory properties of chocolate and its positive effect on human health [12]. Inadequate fermentation of cocoa produces purple and slaty beans, negatively affecting quality and thus marketing price [13]. However, it has been reported that beans without fermentation have genotype-specific sensory attributes, mainly related to bitterness, astringency and acidity, and other attributes depend directly on fermentation, drying and roasting [14].

Cocoa fermentation is mainly carried out by small producers or collection centers in an artisanal manner, with little or no technology and without monitoring of processing conditions, which results in low quality beans [15]. These variations during the fermentation process make the chocolate industry face increasing challenges in maintaining the supply of standard products with high quality organoleptic properties [16]. This condition in the final product causes the cocoa bean to be marketed as ordinary cocoa with prices already regulated by an industry monopolized by large companies, which in the case of Colombia, acquire 90% of the national production [17], which means that the payment received does not have a sufficient impact on the welfare of cocoa-growing families [18], which discourages and puts cocoa production at risk, increasing the possibility of a change in production activity [19].

Although fermentation is an indispensable stage to ensure optimal bean quality [20], there are deficiencies in the number, intensity and coverage of studies on the effects of fermentation and its impact on the physicochemical and sensory properties of the cocoa bean. The objective of this study was to monitor the biochemical, physical and sensory changes during fermentation of cocoa beans in cocoa bean processing plants in the department of Caquetá, Colombia. It is expected that the study will provide a scientific basis for decision-making based on the recognition of the local needs of each plant. With this information, the fermentation processes carried out by cocoa collection centers in the department of Caquetá will be improved and standardized, resulting in better bean quality and added value in marketing.

## 2. Materials and methods

### 2.1. Study area

The study was carried out in the department of Caquetá, specifically in the cocoa processing plants of a) Asociación Orgánica Agrícola de San José del Fragua-ASOACASAN (ASA) (1°19′43″N 75°58′22″W), b) Comité de Cultivadores de Cacao en Sistemas Agroforestales del municipio de San Vicente del Caguán-COMICACAO (CMI) (2°06′55″N 74°46′12″W) and c) Comité de Cacaoteros de los municipios del Paujil y el Doncello-COMCAP (COC) (1°40′26″N 75°16′48″W). The study sites are in an altitudinal range of 280 to 400 m.a.s.l., with an average temperature of 26 °C, relative humidity of 86% and 3,900 mm of precipitation.

### 2.2. Monitoring fermentation and drying of cocoa beans

The cocoa beans were obtained from healthy pods from trees of different universal, national and regional clones obtained from farms associated with the organizations. This process was carried out between March and June 2023, when production peaked. The beans were then fermented and dried according to the protocol of each collection center. The cocoa beans were placed in wooden crates, which were covered with banana leaves and natural jute fiber sacks to maintain a homogeneous temperature. These crates were perforated at the bottom to allow drainage of the cocoa pulp. The fermentation process lasted 168 hours (7 days) and during this time the fermentable mass was turned every 24 hours after 48 hours of fermentation. The turning of the grain was done by passing the dough into the compartments of the fermenter box, a process that was carried out using a plastic shovel. This fermentation process was carried out in triplicate, that is, three fermentation bins were used for each storage center.

During fermentation, the following parameters were monitored daily between 13:00 and 16:00 hours: mass temperature, pH of the grain pulp and pH of the cotyledon. Samples for this monitoring were taken in the center and at the ends of the bin at a depth of 30 cm. This process was carried out in triplicate. The temperature of the dough was determined using a digital thermometer Hl 145 (Hanna Instrument, Woonsocket, RI, USA), the pH of the pulp during fermentation was measured using a pH tester Hl 98108 (Hanna Instrument, Woonsocket, RI, USA) and the pH of the cotyledon was performed following the methodology used by Papalexandratou [21]. For which, 15 g of randomly selected cocoa beans were taken from the three points of the bin; then, the pulp and testa were manually removed from the beans with a knife and the cotyledon was macerated with 30 ml of deionized water in a mortar for one minute until a homogeneous sample was obtained and the pH was read using a pH tester Hl 98108 (Hanna Instrument, Woonsocket, RI, USA).

### 2.3. Determination of chemical characteristics of beans during the fermentation process

#### 2.3.1. Physical analysis

At the end of the fermentation and drying process, a cutting test was carried out according to the NTC 1252 2021 edition [22] to determine the colorations and defects present. For this purpose, 100 cocoa beans were randomly selected from each fermented cocoa pod at each collection center, cut longitudinally with the help of a CocoaT Bean knife (Cocoatown, Alpharetta, Georgia, USA) and the number of beans that were completely fermented, purple, moldy, damaged by insects, and slaty were quantified.

#### 2.3.2. Component Bromatological

Total fat content was determined by the Randall method with petroleum ether (AOAC Method No.991.36) [23], using the semi-automatic solvent extractor (SER 148 VELP Scientific, Italy) whose results were expressed as percentage of fat. Ash content was determined by incinerating the organic matter in a muffle furnace (1,100°C, 22.9A Fisher Scientific, Spain) at 550 ± 5 °C until the sample was free of carbon, cooled in a desiccator and the amount of ash was calculated (AOAC Method No. 930.30) [24], the results were expressed as percentage of ash. Moisture content was determined using the methodology proposed by Nuñez et al. [25] with modifications. A crucible was brought to constant weight, 10 g of dry cocoa beans were weighed into the crucible and left in the oven (Heratherm OMH400, Fisher Scientific, Spain) at 105 ± 5 °C for 4 hours. The results were expressed as percent of moisture. Titratable acidity was measured according to AOAC method 939.05 [24]. Extracts of dried cocoa beans (5 g) were homogenized in 50 mL of deionized water, subsequently filtered through filter paper (3hw, 110 mm, 65 g/m^2^; Boeco, Germany) and titrated with a standard 0.1 N sodium hydroxide (NaOH) solution to a pH of 8.3. The results were expressed as percent of acidity.

#### 2.3.3. Polyphenolic compounds

First, the methanolic extract of defatted cocoa beans was obtained by weighing 3 g and depositing them in falcon tubes, adding 15 mL of methanolic solution (25% HPLC grade methanol, 24% deionized water and 1% HPLC grade formic acid). The samples were homogenized for 25 minutes using vortex and ultrasound for 20 minutes. Subsequently, they were left in darkness for 24 h and centrifuged for 15 min at 4,500 rpm (4 °C) and filtered with filter paper (3hw, 110 mm, 65 g/m^2^; Boeco, Germany). With this extract, the contents of total polyphenols, total flavonoids, DPPH, FRAP and methylxanthine contents were analyzed. The determination of the total polyphenol content was carried out using the Folin-Ciocalteu colorimetric method [26]. Eighteen µL of the extract, 124.5 µL of deionized water, 37.5 µL of Folin-Ciocalteu reagent and 120 µL of 7.1% anhydrous sodium carbonate (Na_2_CO_3_) were taken. It was left to react for 60 min in the dark at room temperature, after which the absorbance was read at 760 nm. Gallic acid was used as a standard. Results were expressed as mg gallic acid equivalent (mg GAE)/g dried cocoa bean. Total flavonoid content was determined by reaction with aluminum chloride (AlCl_3_) according to the methodology proposed by Zhishen et al. [27] with slight modifications. The reaction mixture consisted of 120 µL of deionized water, then 30 µL of the extract was added, followed by 9 µL of 5% sodium nitrite (NaNO_2_) (waited 5 minutes), 9 µL of 10% aluminum chloride (AlCl_3_) (waited 5 minutes), then 60 µL of 1M sodium hydroxide (NaOH) (waited 15 minutes) and finally 72 µL of deionized water. It was left to react in the dark at room temperature for 30 minutes and the absorbance was read at 510 nm. The (+)-catechin was used as a standard for the quantification of total flavonoids. Results were expressed as mg catechin equivalent (mgCE)/g dried cocoa bean.

Quantification of methylxanthines and epicatechin were developed on an Ultimate 3000 HPLC, equipped with an auto-injection system and UV-VIS detector, analytical reverse phase column (Zorbax Eclipse XDB 150mm × 2.1mm) with particle size of 5μm, at 25°C. All compounds were detected at a wavelength of 273 nm. The mobile phase was water/acetic acid (99.7/0.3 v/v) (solvent A) and methanol (Solvent B), flow rate 0.5 mL/min. The gradient was as follows: 0-10 min, 15% linear B; 10.1-18 min, 25% linear B; 18.1-25 min, 30% linear B; 25.1-30 min, 100% linear B; 30.1-35 min, 0 % linear B, followed by 5 min of column re-equilibration before a new injection with an injection volume of 5μL. All analytes were identified and quantified by the external standard method using calibration curves of the standard substances [28].

#### 2.3.4. Antioxidant activity

The DPPH radical scavenging activity (DRSA) was used according to the method of Brand-Williams et al. [29] with slight modifications. A stock solution of DPPH (20 mg/L) was prepared in absolute methanol; the absorbance of the radical was adjusted to 0.3 absorbance units with methanol at 4°C, then 3 µL of the extract and 297 µL of the adjusted DPPH solution were taken. It was allowed to react in the dark for 30 min at room temperature and the absorbance was read at a wavelength of 517 nm. The results were expressed as TEAC values in µmol of Trolox (µmol Trolox)/g of dry cocoa beans, constructing a reference curve using Trolox as an antioxidant. The reducing capacity FRAP (ferric reducing antioxidant power) evaluates the antioxidant capacity of a sample according to its ability to reduce ferric iron (Fe^+3^) present in a complex with 2,4,6-tri(2-pyridyl)-s-triazine (TPTZ), to the ferrous form (Fe^+3^) [30]. The assay was carried out in a pH 3.6 acetic acid-sodium acetate buffer containing TPTZ and FeCl_3_. 15 µL of the extract, 15 µL of buffer and 270 µL of FRAP solution were used as samples.

It was allowed to react in the dark for 30 min at room temperature and the absorbance was read at a wavelength of 590 nm. The FRAP values were expressed as mg ascorbic acid (mg AA/g of dry cocoa bean), based on a reference curve of ascorbic acid as the primary standard.

#### 2.3.5. Sensory analysis

The cocoa paste samples (cocoa bean roasted and milled) were analyzed by a sensory panel following the process described by ICONTEC [22]. The roasting curve was constructed according to bean size and moisture percentage; the roasted beans were also passed through a grinder and the husk was separated from the nibs. Finally, the nibs were processed and refined by a melangeur for the manufacture of cocoa paste/liquor. Sensory evaluation was carried out with the help of five trained panelists, and the main qualities were determined: cocoa, acidity, astringency, bitterness, and notes of fresh fruit, brown, floral, wood, spice and nut, as well as atypical flavors in the samples. The order of perception, intensity, residual flavor and persistence were established as scores given by the panel. The international cocoa evaluation scale of excellence [31] was used, which scores from 0 to 10 points, where 0 indicates the complete absence of the evaluated attribute and 10 indicates a very high intensity.

### 2.4. Data analysis and experimental design

A completely randomized plot design with three replications was used where the site (ASA, CMI and COC) and fermentation time (4 and 7 days) were analyzed on the different variables related to grain physics, bromatological composition, polyphenolic compound content and antioxidant activity as sensory attributes. The data were analyzed by fitting a linear mixed model (LMM) where in the fixed factor the sites (ASA, CMI and COC) and the fermentation time (days 4 and 7) were adjusted, and within the random factor the repetition within the sites. Assumptions of normality and homogeneity of variance were examined through an exploratory analysis of residuals. In addition, the LSD Fisher test (P< 0.05) was performed to determine if differences existed between fixed factors. Line graphs were also made to show the behavior of temperature and pH during fermentation. Subsequently, radar plots were made to visualize the sensory attributes of the sites by days of fermentation. In addition, Pearson’s correlation using the “corrr” package [32] was employed to establish correlations between antioxidant activity, polyphenolic compounds and methylxanthines. Finally, a principal component analysis (PCA) was performed to determine the multivariate relationships between the variables evaluated and the study sites. The models were applied using the statistical software InfoStat [33], PCA and Pearson correlation were performed using the packages ade4, ggplot2 [34], factoextra [35], FactoMineR [36] in the R language software, using the RStudio interface [37].

## 3. Results

### 3.1. Temperature and pH of fermented cocoa beans

During the fermentation period, the temperature increased until 4 days, reaching an average of 45 °C, after which it stabilized (Figure 1a). The pH of the pulp did not vary significantly during the fermentation period, however, there was a small variation at 2 and 3 days between sites (Figure 1b). The pH in the cotyledon at the beginning of the fermentation process ranged from 5.6 to 6.3 with a steady decrease to 4.5 (Figure 1c).

**Figure 1.**
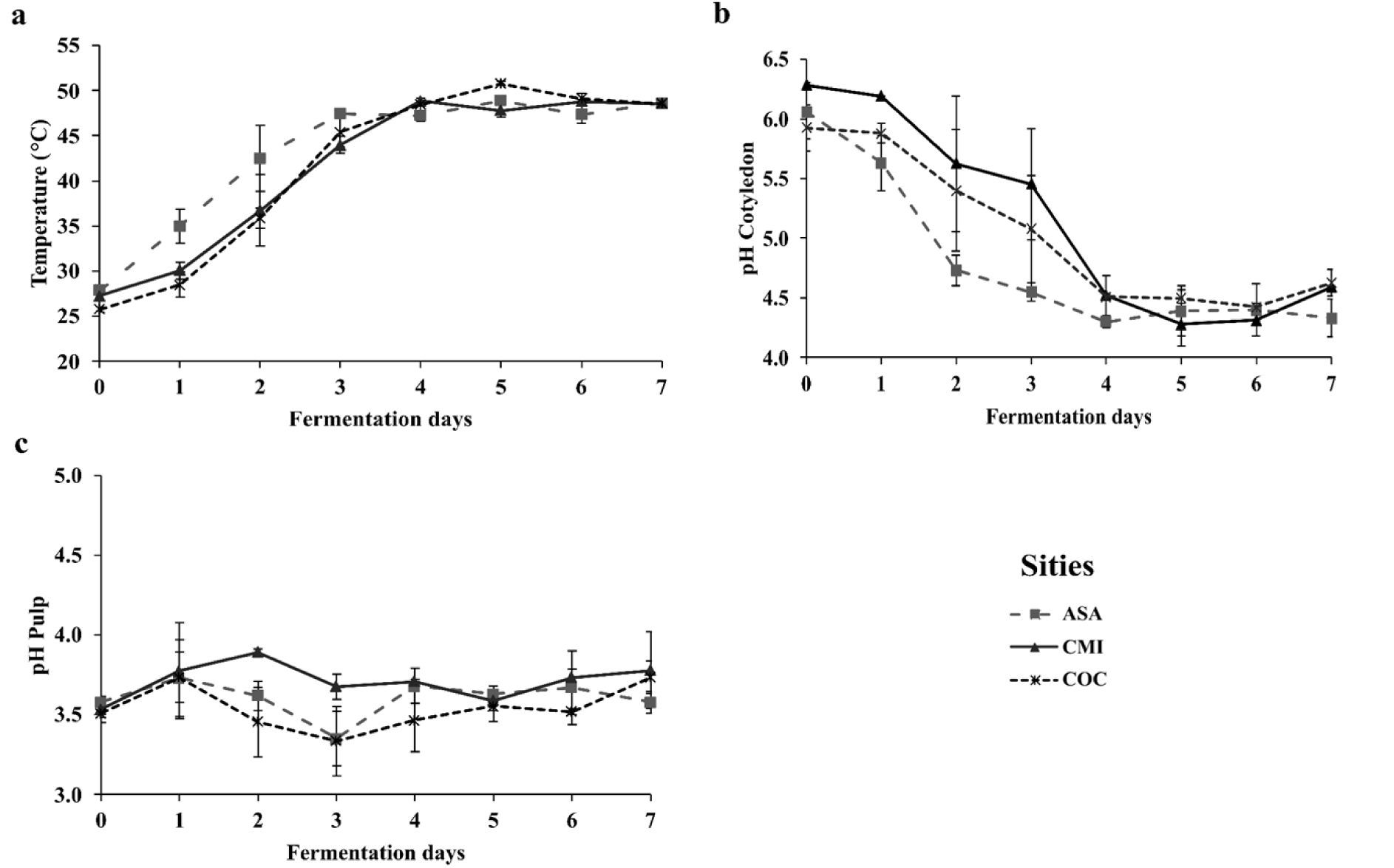
Daily behavior of three variables during cocoa fermentation in three processing plants in the department of Caquetá. ASA: ASOACASAN, CMI: COMICACAO, COC: COMCAP: a) mass temperature, b) pulp pH, c) cotyledon pH. Values correspond to means and standard errors (n=3).

### 3.2. Physical properties of dry cocoa beans

The fermentation process between sites varied (Table 1, P<0.05), a situation that resulted in the number of properly fermented grains; however, no difference was found between fermentation hours. On the contrary, differences between sampling hours were found in the number of violet and partially fermented grains in ASA and totally fermented grains in COC. Between sites, the main difference was in the amount of violet and partially fermented grains (Table 1, P<0.05).

**Table 1.** Results of the shear test on fermented and dried cocoa beans at two fermentation times. Different letters indicate statistically significant differences by LSD Fisher means test (P< 0.05), ^a, b, c^: between the different fermentation hours for each site and ^A, B, C^: between the different sites. ASA: ASOACASAN, CMI: COMICACAO, COC: COMCAP.

| Site | Fermentation time (days) | Fermentation | Fully fermented | Violet | Mouldy beans | Slaty | Partially fermented |
| --- | --- | --- | --- | --- | --- | --- | --- |
| ASA | 4 | 92 ± 2.0 a | 78 ± 2.31 a | 6.67 ± 1.33 b | 0 ± 0.0 a | 1.33 ± 0.67 a | 14 ± 1.15 b |
|  | 7 | 92.7 ± 3.33 a | 73.33 ± 9.4 a | 8.0 ± 4.16 a | 1.33 ± 1.3 a | 0 ± 0.0 a | 17.3 ± 7.06 a |
|  | *General Average* | **92.33 ± 1.74 A** | **75.67 ± 4.45 A** | **7.33 ± 1.98 B** | **0.67 ± 0.67 A** | **0.67 ± 0.42 A** | **15.67 ± 3.28 B** |
| CMI | 4 | 76.7 ± 8.11 a | 58 ± 8.72 b | 21.3 ± 6.36 a | 0.7 ± 0.67 a | 1.33 ± 1.33 a | 18 ± 0.67 b |
|  | 7 | 80 ± 2.0 a | 61.3 ± 2.91 a | 17.3 ± 2.4 a | 1.3 ± 0.67 a | 1.33 ± 0.67 a | 18 ± 1.33 a |
|  | *General Average* | **74.5 ± 3.81 B** | **59.67 ± 4.18 B** | **19.33 ± 3.17 A** | **1.0 ± 0.45 A** | **1.33 ± 0.67 A** | **18.67 ± 0.67 B** |
| COC | 4 | 68 ± 4.0 a | 37 ± 13.0 b | 31 ± 5.0 a | 0 ± 0.0 a | 1 ± 1.0 a | 31 ± 9.0 a |
|  | 7 | 81 ± 9.0 a | 52 ± 2.0 a | 19 ± 9.0 a | 0 ± 0.0 a | 0 ± 0.0 a | 29 ± 7.0 a |
|  | *General Average* | **78.33 ± 5.50 B** | **44.5 ± 6.90 C** | **25.0 ± 5.45 A** | **0.0 ± 0.00 A** | **0.00 ± 0.00 A** | **30.00 ± 4.69 A** |

### 3.3. Component Bromatological

In the different variables of the bromatological component, significant differences were found at the level of fermentation time as well as at the sites (Table 2, P<0.05). For example, fat content and moisture content varied with fermentation time, being significantly higher at seven days; the opposite was true for ash (Table 2, P<0.05). As for the titratable acidity level, no differences were found between fermentation days.

**Table 2.** Bromatological composition of fermented and dried cocoa beans at two fermentation times. Different letters indicate statistically significant differences by LSD Fisher means test (P< 0.05), ^a, b, c^: between the different fermentation hours for each site and ^A, B, C^: between the different sites. ASA: ASOACASAN, CMI: COMICACAO, COC: COMCAP.

| Site | Fermentation time (days) | Fat | Ash | Moisture | Acidity titratable |
| --- | --- | --- | --- | --- | --- |
| ASA | 4 | 40.20 ± 0.28 b | 2.77 ± 0.003 a | 4.09 ± 0.01 a | 0.40 ± 0.02 a |
|  | 7 | 42.49 ± 0.13 a | 2.78 ± 0.003 a | 3.75 ± 0.02 b | 0.41 ± 0.003 a |
| *General Average* |  | **41.34 ± 0.53 B** | **2.77 ± 0.003 A** | **3.98 ± 0.8 A** | **0.40 ± 0.01 C** |
| CMI | 4 | 43.68 ± 0.06 b | 2.84 ± 0.05 a | 3.16 ± 0.03 a | 0.47 ± 0.01 a |
|  | 7 | 45.47 ± 0.34 a | 2.48 ± 0.01 b | 3.31 ± 0.14 a | 0.48 ± 0.01 a |
| *General Average* |  | **44.58 ± 0.43 A** | **2.66 ± 0.8 A** | **3.23 ± 0.07 B** | **0.48 ± 0.01 B** |
| COC | 4 | 42.55 ± 0.74 a | 2.75 ± 0.05 a | 3.21 ± 0.02 a | 0.52 ± 0.003 a |
|  | 7 | 43.74 ± 0.44 b | 2.51 ± 0.01 b | 3.08 ± 0.01 b | 0.55 ± 0.01 a |
| *General Average* |  | **43.15 ± 0.47 A** | **2.63 ± 0.06 A** | **3.15 ± 0.03 B** | **0.53 ± 0.01 A** |

### 3.4. Polyhenolic compounds

At the site level, TPC, TF, DPPH and FRAP contents were statistically different, with COC being different from the other sites (P<0.05, Figure 2). Total polyphenol content was higher at the COC site (507.05 mg GAE/g Cocoa Figure 2a) with respect to the other sites (< 360 mg GAE/g Cocoa). The TF content followed a similar behavior to TPC, with significant differences between sites (P<0.05) and differences between fermentation times for ASA. The TF was higher in COC (309.1 mg Catechin/g cocoa, Figure 2b) with respect to CMI (215.59 mg Catechin/g cocoa) and ASA (185.7 mg Catechin/g cocoa). Values in DPPH (Figure 2c) ranged from 5869.3 to 7781.8 𝜇mol Trolox/g cocoa and for the FRAP assay ranged from 369.8 to 606.7 mg AA/g cocoa among the sites (Figure 2d). At the methylxanthine level, theobromine was different between sites and only different at the level of hours of fermentation in COC (P<0.05, Figure 2e). In the case of caffeine and epicatechin, no differences were found between sites (Figure 2fg); however, only for caffeine there were differences between the fermentation hours for the three sites, while for epicatechin only COC showed differences in fermentation hours.

**Figure 2.**
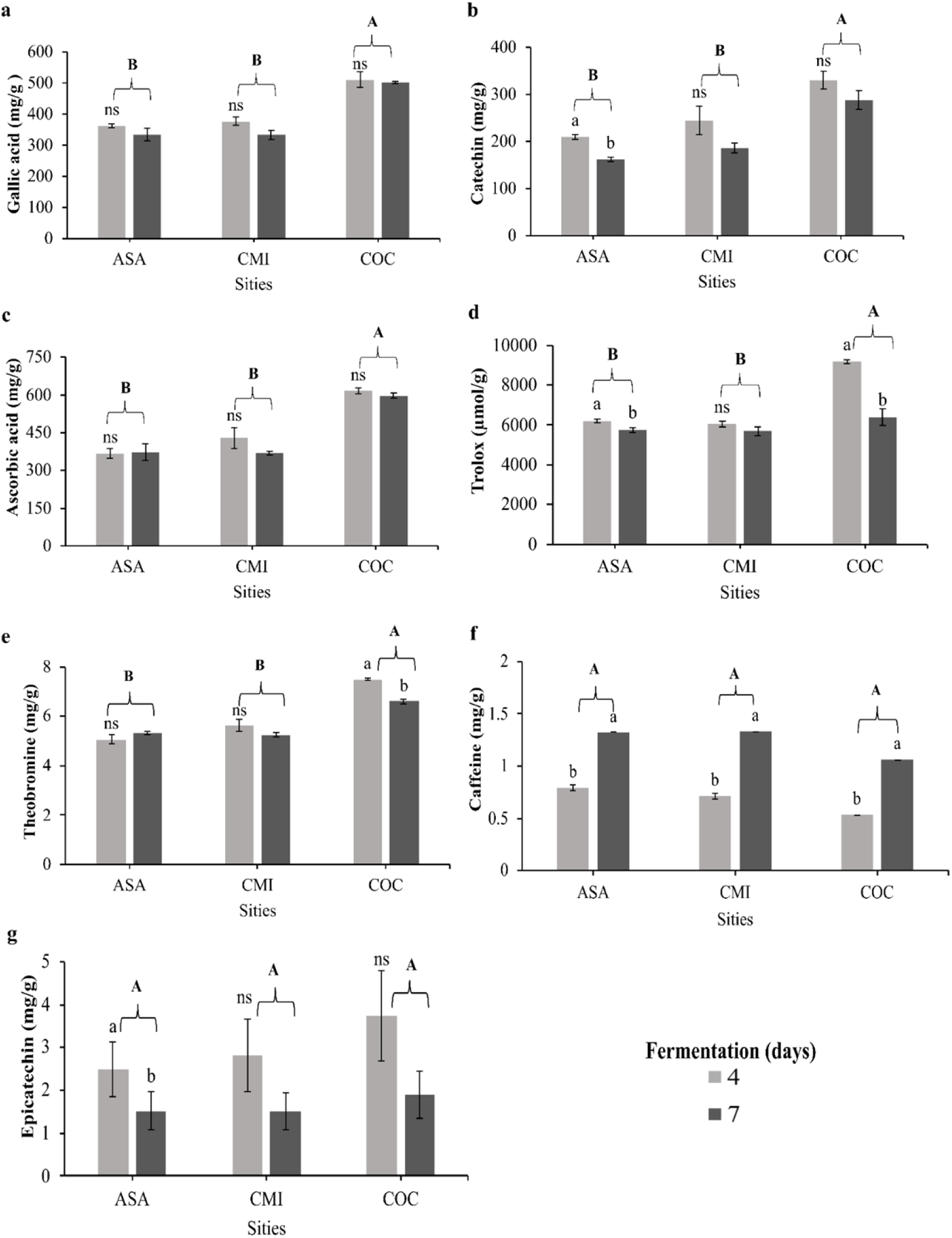
Content of polyphenols, methylxanthines and antioxidant activity in fermented cocoa beans. Different letters indicate statistically significant differences by LSD Fisher means test (P< 0.05), ^a, b, c^: between the different fermentation hours for each site and ^A, B, C^: between the different sites. ASA: ASOACASAN, CMI: COMICACAO, COC: COMCAP.

### 3.5. Sensory profile

In general, the presence of complementary attributes such as nutty, spicy, woody, floral and fresh fruit was found at all sites (Figure 3). It was evident that COC presented higher acidity (5.5) with respect to ASA (3.67) and CMI (3.33) on the fourth day of fermentation; however, on day 7 acidity decreased in COC (4.5) but increased in CMI (5.0). Astringency and bitterness increased with days of fermentation in ASA and COC, while astringency and bitterness decreased with the course of fermentation in CMI.

**Figure 3.**
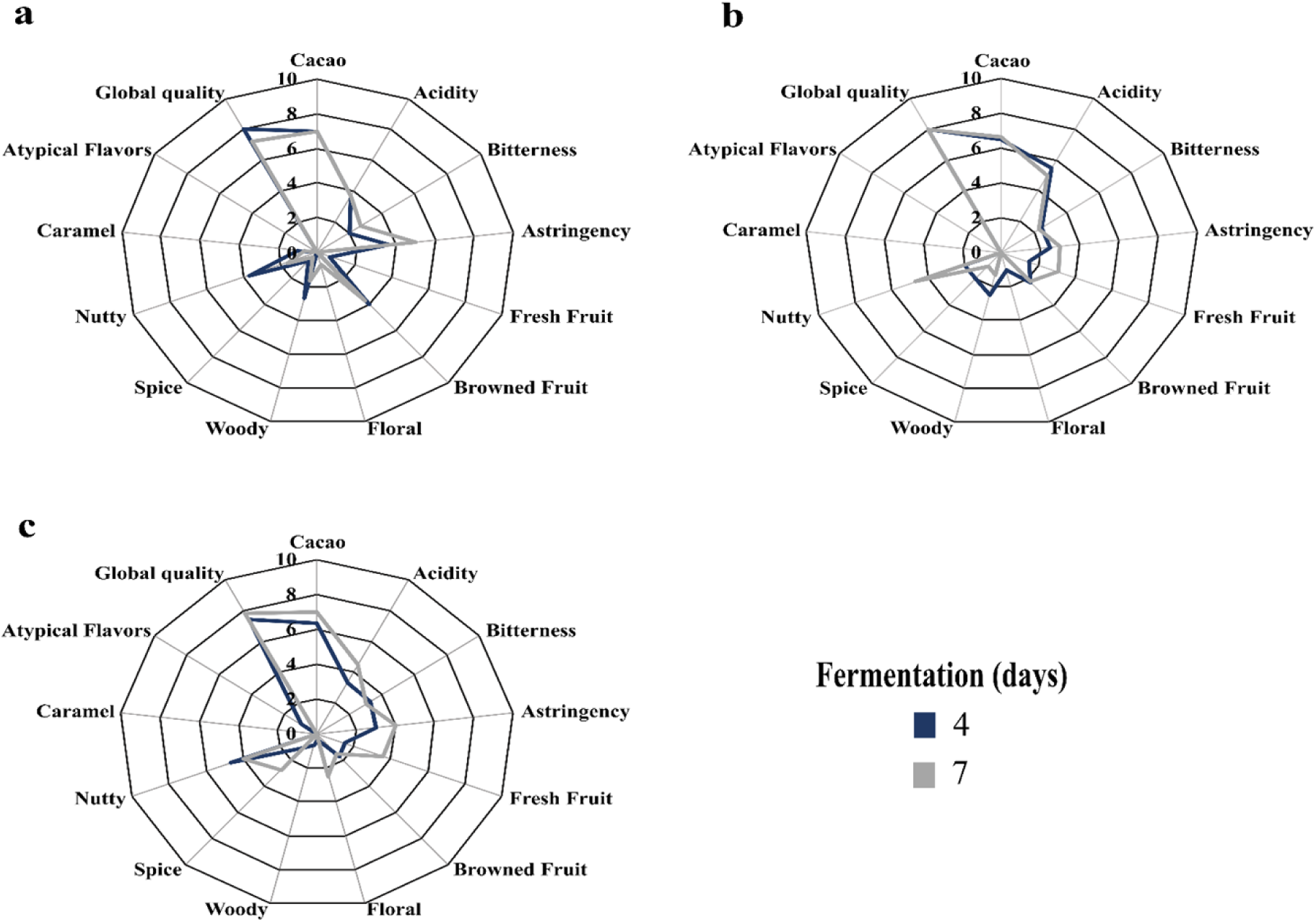
Sensory profile of cocoa liquor in three processing plants: a. ASA: ASOACASAN, b. CMI: COMICACAO, c. COC: COMCAP.

Figure 4 shows the different positive and negative correlations between the different variables analyzed. A positive correlation was found between polyphenolic compounds and antioxidants with theobromine. Likewise, titratable acidity had a positive correlation with total polyphenols, total flavonoids and variables related to antioxidant capacity (FRAP and DPPH). On the other hand, caffeine was negatively correlated with DPPH, theobromine and epicatechin (Figure 4).

**Figure 4.**
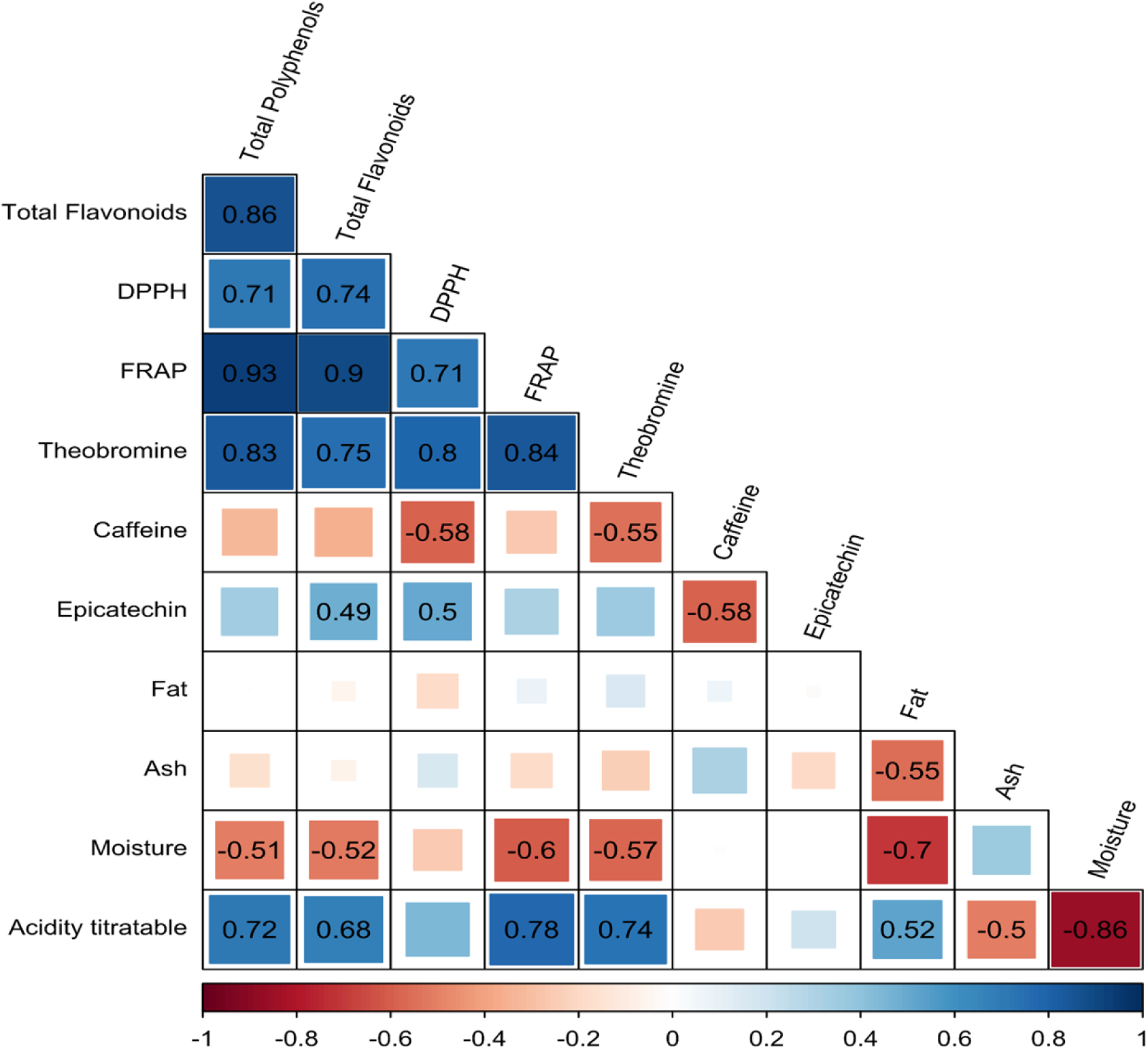
Correlation analysis between the different variables analyzed. The numbers shown presented significant correlations (P<0.05) in the color gradient from red to blue showing negative and positive correlation, respectively.

Principal component analysis explained 43.8% of the variance (PCA, Figure 5), with CP1 explaining 27.2% and opposing the ASA and COC sites by variables such as moisture, titratable acidity as well as phenolic compound content and antioxidant activity. CP2 explained 16.6%, opposing different sensory attributes such as cocoa and bitterness. According to the Monte-Carlo test, the sites explained 24.9% of the variance (Figure 5).

**Figure 5.**
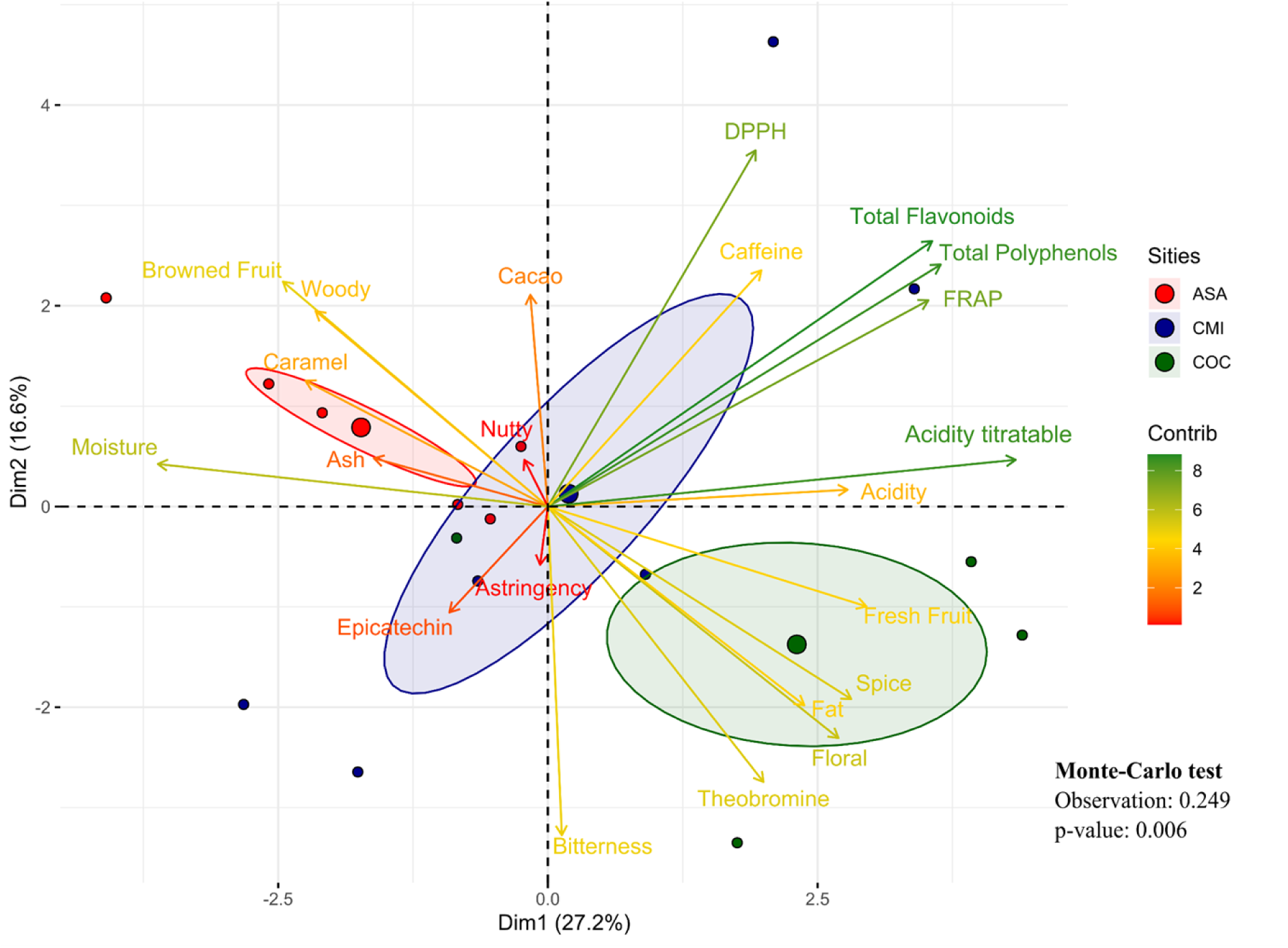
PCA projection of variables related to bromatological composition, polyphenol content, methylxanthines and antioxidant activity in cocoa beans fermented in different cocoa bean processing plants in the department of Caquetá, Colombia. ASA: ASOACASAN, CMI: COMICACAO, COC: COMCAP. Contribution of each of the variables evaluated in the PC1/PC2 principal components of PCA, the gradient from green to red means from greater to lesser contribution.

## 4. Discussion

### 4.1. Temperature and pH of fermented cocoa beans

An increase in temperature was found in the fermentation mass due to the activity carried out by the yeasts involved in the metabolization of sugars in the cocoa pulp (exothermic process), to produce ethanol and other metabolites (other alcohols, esters, aldehydes and ketones), a process that releases heat and increases the fermentation temperature [38]. It was observed that from the fifth day of fermentation the temperature did not vary, reaching a temperature between 40-50 °C, a desirable range for good fermentation [39]. On the other hand, from the third day of fermentation the mass recorded an increase in pH, a situation attributed to the decrease by leaching of citric acid contained in the pulp [40] and the decrease of volatile organic acids leading to an increase in pH in the fermentation mass [41] as to the metabolization of citric acid by the yeasts [42]. This decrease in pH within the first few days at all sites is attributed mainly to the diffusion of organic acids in the bean produced by lactic acid bacteria [43] and together with the high temperatures, which lead to embryo death, triggering a series of biochemical changes that greatly impact the development of bean flavor and color. Also, this behavior was considered [44] to be because of turning the cocoa mass after 48 hours, because it favors aeration and the growth of acid-acetic bacteria [45]. Finally, the pH of the cotyledon at the end of fermentation in the three sites was very similar ASA (4.33), CMI (4.59) and COC (4.63), these results are very important since some studies mention that pH above 5.5 at the end of fermentation indicates lower quality fermented beans [46], however, Calvo et al. (2021) [16] mentions that pH ranges in cotyledon between 4.8 and 5.2 indicates a good fermentation process. In our study, fermentation at CMI and COC were the closest to the conditions.

### 4.2. Physical properties of dry cocoa beans

Variation was found in the number of fermented grains according to the category. For example, the increase in the percentage of fermented and fully fermented beans (brown) is mainly associated with anthocyanin degradation [47] because of gradually increasing temperature in the fermentation box [48]. Purple cocoa beans are characteristic of an inadequately fermented bean and brown beans are typical of optimal fermentation [49], a situation that was present in the CMI and COC sites. The percentage of partially fermented beans presented significant differences between sites, with higher values in COC. The presence of partially fermented beans is produced by some factors such as: reduction in the amount of turning in the cocoa mass during fermentation or due to the location of the beans in the fermentation box [50]. For this reason, several studies mention the importance and impact of turning on the physical characteristics of cocoa beans [51].

### 4.3. Component Bromatological

An increase in fat content was evidenced with increasing days of fermentation at all sites (P<0.05), the data obtained agree with the study of Millena et al. [52] who found an increase in fat content from the fifth day of fermentation. The increase in fat contents during fermentation days can be attributed to the interaction between acidity and temperature during fermentation, leading to the alteration of lipid bonds [53]. On the other hand, the values in ash percentage were below 3%, values that coincide with those reported by Calvo et al. [16], likewise, this decrease in ash content during fermentation was reported by Afoakwa et al. [54]. This behavior is explained by the loss of water-soluble minerals drained during the fermentation process [55]. Moisture contents varied significantly in all sites, being ASA with the highest values (3.98%) with respect to CMI and COC, it is believed that this behavior is due to the geographical location and precipitation of the site, because ASA is located in the Andean-Amazonian transition with higher precipitation than the other sites included in the study; according to Bomdzele and Molua [56] high precipitation can slow grain drying and increase moisture content. The decrease in grain moisture content during fermentation at the COC site is explained by the increase in temperature allowing moisture to diffuse outward and by the effect of turning the fermentable mass [57]. The increase in acidity percentage is associated with the increase of lactic acid and acetic acid during the fermentation days [16]. These acids enter the cotyledon generating biochemical reactions and causing embryo death [58].

### 4.4. Polyphenolic compounds

Total polyphenol content was higher at the COC site (507.05 mg GAE/g cocoa) compared to the other sites (< 360.00 mg GAE/g cocoa), the highest COC contents are mainly attributed to beans from hybrid cocoa plantations found in the study area. According to Jonfia-Essien et al. [59] hybrid cocoa beans have higher polyphenol levels and antioxidant activity than clones, due to their genetic variability. Regarding total polyphenol values, our results were much higher than those reported by Ramón et al. (2022) [4] in fermented and roasted beans (210.2 mg GAE/g cocoa), with the difference that in our study, the cocoa beans evaluated were not roasted, only fermented and dried. According to Urbańska et al. [60] the polyphenol content may decrease due to the oxidation of these compounds by the effect of high temperatures in the roasting process and interaction with Maillard reaction products [61,62]. In addition to the above, a decrease in total polyphenol content was observed at all sites with increasing fermentation time, which is attributed to the effect of increasing temperature in the fermentation box [16], although this decrease has also been reported due to the influence of polyphenol oxidase enzyme activity during fermentation and drying [63]. In general terms, we posit that grains that have undergone adequate fermentation have lower polyphenol contents. This is desirable to achieve pleasant sensory profiles, because higher polyphenols content are associated with astringency [64], while anthocyanins are linked to the violet color of unfermented grains [65]. Both aspects are considered undesirable in chocolate production [66,67]. However, their decrease has a proportional effect on antioxidant activity, a desirable feature from the functional point of view of chocolate to decrease the risk of oxidative stress-mediated diseases [68].

Total flavonoids content (TF) followed a similar behavior that TPC, with significant differences between sites (P<0.05) and differences between fermentation times for ASA. The TF was higher in COC (309.1 mg Catechin/g cocoa) with respect to CMI (215.6 mg Catechin/g cocoa) and ASA (185.7 mg Catechin/g cocoa). Our results showed changes in TF contents because of fermentation days, this explains that flavonoids also decreased during fermentation as did polyphenols, corroborating what was found by Hu et al. [69] in their research on chemical properties in roasted and unroasted cocoa beans, in which they mention that flavonoids are more sensitive to high temperatures than polyphenols. The values found in TF in our study were higher than those reported by da Silva Oliveira et al. [70] and like those found by Cuéllar-Álvarez et al. [71] in copoazú (*Theobroma grandiflorum*) beans with different fermentation days. The literature primarily reports the total flavonoid content extracted from fermented and roasted grains; hence, our values are higher than those reported by many authors, because in our study the grains used for physical and chemical properties analysis did not go through the roasting process. Zaman et al. [72] mentions that roasting causes the contents of flavonoids, flavonols, flavones and anthocyanins to decrease with increasing time and temperature during roasting. While flavonoids are important bioactive compounds that have been associated with their anti-inflammatory properties, they have a negative effect on the sensory attributes of the bean; however, chocolate manufacturers today pay great attention to declaring their products as functional foods [73]. This opens the door to discussions regarding the manufacture of chocolates with fermented/roasted beans, fermented/unroasted beans or fermented/roasted beans enriched with encapsulated polyphenols.

### 4.5. Sensory profile

In general, complementary attributes such as nutty, spicy, woody, floral and fresh fruit were present in all sites, which are categorized as fine cocoa aroma and flavor [74]. The sensory evaluation revealed that, in all sites and days of fermentation, cocoa notes obtained intensity scores higher than 6, however, our results contrast with those reported by Calvo et al. [16] who found values below 5 in cocoa samples from three different regions in Colombia. Also, Romero and Pabón [75] mentions that beans with high cocoa flavor intensity indicate a successful processing (fermentation and drying), although the main attributes are mostly influenced by genotype [76]. In fermentation and drying, pyrazine contents increase because they are compounds of thermal origin and increase with increasing temperature [77], some studies mention that pyrazines are related to cocoa flavor [8].

It was evidenced that COC presented higher acidity (5.5) with respect to ASA (3.7) and CMI (3.3) on the fourth day of fermentation, however, on day 7 acidity decreased in COC (4.5), but increased in CMI (5.0), this decrease in acidity is mainly due to acetic acid volatilization during fermentation and drying because of temperature [78]. The increase in acidity during fermentation at CMI was mainly attributed to the presence of volatile acids, generated as a product of over fermentation and poor drying [54,79]. Acidity is an important attribute, as high levels of acidity in cocoa samples can be detrimental to bean quality [80]. Astringency and bitterness increased with fermentation days in ASA and COC, this behavior was associated with the effect of a possible over fermentation, we also believe that the increase in astringency is due to a deficiency in the turnovers of the fermenting mass [81,82] behavior that contrasts with a good fermentation [83] while astringency and bitterness decreased with the course of fermentation at CMI, this decrease is characteristic of a good fermentation process [84].

This flavor is acquired mainly in the roasting process, because it has a direct effect on the development of flavors in the cocoa beans through various reactions, including Maillard reactions, which contribute to the presence of nutty flavor and aroma [85]. Rodriguez-Campos et al. [80] concluded that 6 days of fermentation are sufficient to produce volatile compounds with desirable flavor notes in cocoa, conclusions that contrast with the results of this study, given that the profiles obtained here had flavor notes characteristic of over fermentation in ASA, although favorable notes in COC and CMI with 7 days of fermentation. In addition, it is known that cocoa flavor and complementary attributes are influenced by the genetic potential of cocoa, but also by postharvest practices (fermentation and drying) [86].

## 5. Conclusions

The management of the fermentation process has a significant impact on the different characteristics (biochemical, physical and sensory) of the cocoa beans. A similar trend of fermentation mass variables was found at all sites where cotyledon pH decreased during the fermentation process and fat and moisture content varied with fermentation time. At the site level, total polyphenol (TPC), total flavonoids (TF), DPPH and FRAP contents were statistically different, with COC being different from the other sites. The TPC was higher in the COC site (507 mg GAE/g Cocoa) with respect to the other sites (< 360 mg GAE/g Cocoa). The TF content followed a similar behavior to TPC, with significant differences between sites and differences between fermentation times for ASA. The TF was higher in COC (309.1 mg Catechin/g) with respect to CMI (215.6 mg Catechin/g) and ASA (185.7 mg Catechin/g). Values in DPPH ranged from 5869.3 to 7781.8 𝜇mol Trolox/g cocoa and for the FRAP assay ranged from 369.8 to 606.7 mg AA/g cocoa among sites. Complementary attributes such as nutty, spicy, woody, woody, floral and fresh fruit were observed at all sites. Astringency and bitterness increased with days of fermentation at ASA and COC, while astringency and bitterness decreased with the course of fermentation at CMI.

## Acknowledgements

We are grateful to the Minciencias project "Fortalecimiento de vocaciones científicas en jóvenes a través de becas de pasantía en la región centro sur: Caquetá, Amazonas, Putumayo, Huila, Tolima" financed with resources from the general royalty’s system and executed by the Vice-Rectory of Research and Innovation of the Universidad de la Amazonia. We thank the University of Amazon for supporting this study through the strategy of hiring research assistants in the research support units at the CIMAZ Macagual Amazon Research Center in the framework of the "Pro-development of the University" stamp of the Amazon" law 1301 of 2009. We also thank the following collaborators: Asociación Orgánica Agrícola de San José del Fragua (ASOACASAN), Comité de Cultivadores de Cacao en Sistemas Agroforestales del municipio de San Vicente del Caguán (COMICACAO) and Comité de Cacaoteros de los municipios del Paujil y el Doncello (COMCAP) for providing the benefit facilities to carry out this study. Finally, we would like to thank Corporacion Econexus Colombia (INSITU) for their support in the tasting of the samples.

## References

1. Goya L, Kongor JE, de Pascual-Teresa S. From Cocoa to Chocolate: Effect of Processing on Flavanols and Methylxanthines and Their Mechanisms of Action. International Journal of Molecular Sciences 2022, Vol 23, Page 14365. 2022;23: 14365. doi:10.3390/IJMS232214365

2. Gavrilova NG. Contemporary global production and consumption of cocoa: An assessment. IOP Conf Ser Earth Environ Sci. 2021;839. doi:10.1088/1755-1315/839/2/022095

3. Hirko B, Mitiku H, Getu A. Role of fermentation and microbes in cacao fermentation and their impact on cacao quality. Systems Microbiology and Biomanufacturing 2023. 2023;1: 1–12. doi:10.1007/S43393-023-00160-9

4. Ramón V, Hernández HE, Polania P, Suárez JC. Spatial Distribution of Cocoa Quality: Relationship between Physicochemical, Functional and Sensory Attributes of Clones from Southern Colombia. Agronomy 2023, Vol 13, Page 15. 2022;13: 15. doi:10.3390/AGRONOMY13010015

5. Pallares-Pallares A, Perea-Villamil JA, Javier López-Giraldo L. Impacto de las condiciones de beneficio sobre los compuestos precursores de aroma en granos de cacao (Theobroma cacao L) del clon CCN-51. Respuestas, ISSN 0122-820X, ISSN-e 2422-5053, Vol 21, N° 1, 2016, págs 120-133. 2016;21: 120–133. Available: https://dialnet.unirioja.es/servlet/articulo?codigo=5598206&info=resumen&idioma=S PA

6. Polanía-Hincapié PA, Suárez JC, Hernández HE, Ramón-Triana VY, Cuéllar-Álvarez LN, Casanoves F. Influence of Fermentation Time on the Chemical and Functional Composition of Different Cocoa Clones from Southern Colombia. Fermentation 2023, Vol 9, Page 982. 2023;9: 982. doi:10.3390/FERMENTATION9110982

7. Niikoi Kotey R, Asomaning Odoom D, Kumah P, Oppong Akowuah J, Fobi Donkor E, Kwatei Quartey E, et al. Effects of Fermentation Periods and Drying Methods on Postharvest Quality of Cocoa (Theobroma Cacao) Beans in Ghana. J Food Qual. 2022;2022. doi:10.1155/2022/7871543

8. Sari ABT, Fahrurrozi, Marwati T, Djaafar TF, Hatmi RU, Purwaningsih, et al. Chemical Composition and Sensory Profiles of Fermented Cocoa Beans Obtained from Various Regions of Indonesia. Int J Food Sci. 2023;2023. doi:10.1155/2023/5639081

9. García-Rincón PA, Núñez-Ramírez JM, Bahamón-Monje AF, García-Rincón PA, Núñez-Ramírez JM, Bahamón-Monje AF. Physicochemical and sensory characteristics of fermented almonds of national cacao (Theobroma Cacao L.) with addition of probiotics in the amazonic research center, Cimaz Macagual (Caquetá, Colombia). Ingeniería y competitividad. 2021;23. doi:10.25100/iyc.v23i2.10885

10. De Vuyst L, Leroy F. Functional role of yeasts, lactic acid bacteria and acetic acid bacteria in cocoa fermentation processes. FEMS Microbiol Rev. 2020;44: 432–453. doi:10.1093/femsre/fuaa014

11. Loo-Miranda JLM, Chire-Fajardo GС, Ureña-Peralta MO, Loo-Miranda JLM, Chire-Fajardo GС, Ureña-Peralta MO. Correlation between Electrical Conductivity and the Percentage of Fermented Beans for Peruvian CCN 51 Cocoa Beans. Ingeniería e Investigación. 2022;42. doi:10.15446/ING.INVESTIG.92556

12. Soares TF, Oliveira MBPP. Cocoa by-products: characterization of bioactive compounds and beneficial health effects. Molecules. 2022;27: 1625.

13. Ackah E, Dompey E. Effects of fermentation and drying durations on the quality of cocoa (Theobroma cacao L.) beans during the rainy season in the Juaboso District of the Western-North Region, Ghana. Bull Natl Res Cent. 2021;45: 175. doi:10.1186/s42269-021-00634-7

14. Alvarez-Villagomez KG, Ledesma-Escobar CA, Priego-Capote F, Robles-Olvera VJ, García-Alamilla P. Influence of the starter culture on the volatile profile of processed cocoa beans by gas chromatography–mass spectrometry in high resolution mode. Food Biosci. 2022;47: 101669. doi:10.1016/J.FBIO.2022.101669

15. Forte M, Currò S, Van de Walle D, Dewettinck K, Mirisola M, Fasolato L, et al. Quality Evaluation of Fair-Trade Cocoa Beans from Different Origins Using Portable Near-Infrared Spectroscopy (NIRS). Foods. 2023;12. doi:10.3390/FOODS12010004/S1

16. Calvo AM, Botina BL, García MC, Cardona WA, Montenegro AC, Criollo J. Dynamics of cocoa fermentation and its effect on quality. Scientific Reports 2021 11:1. 2021;11: 1–15. doi:10.1038/s41598-021-95703-2

17. Ramírez Chamorro LE, Abaunza González CA, Rodríguez Polanco L, Varón Devia EH, Barragán Quijano E, Rojas-Molina J. Modelo productivo para el cultivo de cacao (Theobroma cacao) para el departamento del Huila. AGROSAVIA. Mosquera, Colombia: AGROSAVIA; 2020. Available: http://editorial.agrosavia.co/index.php/publicaciones/catalog/download/29/20/361-1?inline=1

18. Jaimes Suárez YY, Castañeda GAA, Daza EYB, Bustos FM, Silva RAC, Estrada GAR, et al. Modelo productivo para el cultivo de cacao (Theobroma cacao L.) en el departamento de Boyacá. Editorial AGROSAVIA. 2022. doi:10.21930/AGROSAVIA.MODEL.7405590

19. Hashmiu I, Agbenyega O, Dawoe E. Determinants of crop choice decisions under risk: A case study on the revival of cocoa farming in the Forest-Savannah transition zone of Ghana. Land use policy. 2022;114: 105958. doi:10.1016/j.landusepol.2021.105958

20. De Vuyst L, Weckx S. The cocoa bean fermentation process: from ecosystem analysis to starter culture development. J Appl Microbiol. 2016;121: 5–17. doi:10.1111/jam.13045

21. Papalexandratou Z, Falony G, Romanens E, Jimenez JC, Amores F, Daniel HM, et al. Species Diversity, Community Dynamics, and Metabolite Kinetics of the Microbiota Associated with Traditional Ecuadorian Spontaneous Cocoa Bean Fermentations. Appl Environ Microbiol. 2011;77: 7698. doi:10.1128/AEM.05523-11

22. INCONTEC. NTC-ISO 2292:2021 Cacao en grano. Muestreo. 2021.

23. AOAC. Official Method 991.36, Fat (crude) in meat, solvent extraction (Submersion) method. Off. Methods Anal. Arlington, VA; 1995.

24. AOAC. International A: Official Methods of Analysis of the AOAC International. The Association: Arlington County, VA, USA. 2000.

25. Nuñez JM, Bahamón Monje AF, García Rincón PA. Características fisicoquímicas y sensoriales de almendras fermentadas de cacao nacional (Theobroma Cacao L.) con adición de probióticos en el centro de investigaciones amazónicas, Cimaz Macagual (Caquetá, Colombia). Universidad del Valle; 2021. doi:10.25100/iyc.v23i2.10885

26. Singleton VL, Rossi JA. Colorimetry of total phenolics with phosphomolybdic-phosphotungstic acid reagents. Am J Enol Vitic. 1965;16: 144–158.

27. Zhishen J, Mengcheng T, Jianming W. The determination of flavonoid contents in mulberry and their scavenging effects on superoxide radicals. Food Chem. 1999;64: 555–559.

28. Quiroga Ruiz Y, Herrera Sánchez DA. Efecto de la adición de polifenoles sobre las características químicas y sensoriales de un chocolate. Thesis, Universidad Industrial de Santander. 2019. Available: https://noesis.uis.edu.co/handle/20.500.14071/13780

29. Brand-Williams W, Cuvelier ME, Berset C. Use of a free radical method to evaluate antioxidant activity. LWT - Food Science and Technology. 1995;28: 25–30. doi:10.1016/S0023-6438(95)80008-5

30. Benzie IFF, Strain JJ. The ferric reducing ability of plasma (FRAP) as a measure of “antioxidant power”: the FRAP assay. Anal Biochem. 1996;239: 70–76. doi:10.1006/ABIO.1996.0292

31. COEX. Home: International Standards for the Assessment of Cocoa Quality and Flavour. Brigitte Laliberté DANV (Bioversity I and SF (RB-E, editor. 2023. Available: https://www.cocoaqualitystandards.org/

32. Kuhn M, Jackson S, Cimentada J. corrr: Correlations in R. R package version 04. 2020;3: 1–15.

33. Di Rienzo J, Balzarini M, Gonzalez L, Casanoves F, Tablada M, Walter Robledo C. Infostat: software para análisis estadístico. 2010.

34. Wickham H. Programming with ggplot2. 2016; 241–253. doi:10.1007/978-3-319-24277-4_12

35. Kassambara A, Mundt F. Package ‘factoextra.’ Extract and visualize the results of multivariate data analyses. 2017;76.

36. Husson F, Josse J, Le S, Mazet J, Husson MF. Package ‘factominer.’ 2008;96: 698.

37. RStudio Team. RStudio: Integrated Development Environment for R. RStudio, PBC; 2022. Available: http://www.rstudio.com/

38. Cortez D, Quispe-Sanchez L, Mestanza M, Oliva-Cruz M, Yoplac I, Torres C, et al. Changes in bioactive compounds during fermentation of cocoa (Theobroma cacao) harvested in Amazonas-Peru. Curr Res Food Sci. 2023;6: 100494. doi:10.1016/J.CRFS.2023.100494

39. Dewandari KT, Rahmawati R, Munarso SJ. The effect of techniques and fermentation time on cocoa beans quality (Theobroma cacao L.). IOP Conf Ser Earth Environ Sci. 2021;653: 012046. doi:10.1088/1755-1315/653/1/012046

40. Peña González MA, Ortiz Urgiles JP, Santander Pérez FA, Lazo Vélez MA, Caroca Cáceres RS. Physicochemical changes during controlled laboratory fermentation of cocoa (CCN-51) with the inclusion of fruits and on-farm inoculation. Brazilian Journal of Food Technology. 2023;26: e2023013. doi:10.1590/1981-6723.01323

41. Van de Voorde D, Díaz-Muñoz C, Hernandez CE, Weckx S, De Vuyst L. Yeast strains do have an impact on the production of cured cocoa beans, as assessed with Costa Rican Trinitario cocoa fermentation processes and chocolates thereof. Front Microbiol. 2023;14: 1232323. doi:10.3389/FMICB.2023.1232323/BIBTEX

42. Alvarez JP. AROMA-PRODUCING YEASTS ASSOCIATED WITH COCOA BEANS FERMENTATION: STARTER CULTURE SELECTION FOR FLAVOR MODULATION OF CHOCOLATE. Thesis, Universidade Federal do Tocantins . 2017.

43. Korcari D, Fanton A, Ricci G, Rabitti NS, Laureati M, Hogenboom J, et al. Fine Cocoa Fermentation with Selected Lactic Acid Bacteria: Fermentation Performance and Impact on Chocolate Composition and Sensory Properties. Foods. 2023;12. doi:10.3390/FOODS12020340/S1

44. Díaz-Muñoz C, De Vuyst L. Functional yeast starter cultures for cocoa fermentation. J Appl Microbiol. 2022;133: 39–66. doi:10.1111/JAM.15312

45. Deus VL, Bispo ES, Franca AS, Gloria MBA. Understanding amino acids and bioactive amines changes during on-farm cocoa fermentation. Journal of Food Composition and Analysis. 2021;97: 103776. doi:10.1016/J.JFCA.2020.103776

46. Lee AH, Neilson AP, O’Keefe SF, Ogejo JA, Huang H, Ponder M, et al. A laboratory-scale model cocoa fermentation using dried, unfermented beans and artificial pulp can simulate the microbial and chemical changes of on-farm cocoa fermentation. European Food Research and Technology. 2019;245: 511–519. doi:10.1007/S00217-018-3171-8/FIGURES/5

47. López-Hernández M del P, Melo-Martinez SE, Criollo-Núñez J. Effect of the maturity stage, genotype, and geographical location on the physicochemical characteristics of the cocoa bean during fermentation. INGENIERÍA Y COMPETITIVIDAD. 2023;25. doi:10.25100/IYC.V25I2.12503

48. Delgado JD, Mandujano JI, Reátegui D, Ordoñez ES. Desarrollo de chocolate oscuro con nibs de cacao fermentado y no fermentado: polifenoles totales, antocianinas, capacidad antioxidante y evaluación sensorial. Scientia Agropecuaria. 2018;9: 543–550. doi:10.17268/SCI.AGROPECU.2018.04.10

49. Hernández-Hernández C, López-Andrade PA, Ramírez-Guillermo MA, Guerra Ramírez D, Caballero Pérez JF. Evaluation of different fermentation processes for use by small cocoa growers in mexico. Food Sci Nutr. 2016;4: 690–695. doi:10.1002/fsn3.333

50. Guzmán-Alvarez RE, Márquez-Ramos JG, Guzmán-Alvarez RE, Márquez-Ramos JG. Fermentation of Cocoa Beans. Fermentation - Processes, Benefits and Risks. 2021 [cited 23 Aug 2023]. doi:10.5772/INTECHOPEN.98756

51. Hamdouche Y, Meile JC, Lebrun M, Guehi T, Boulanger R, Teyssier C, et al. Impact of turning, pod storage and fermentation time on microbial ecology and volatile composition of cocoa beans. Food Research International. 2019;119: 477–491. doi:10.1016/j.foodres.2019.01.001

52. Millena CG, Balonzo ARR, Rentoy JR, Ruivivar SS, Bobiles SC. Effect of fermentation stages on the nutritional and mineral bioavailability of cacao beans (Theobroma cacao L.). Journal of Food Composition and Analysis. 2023;115: 104886.

53. Servent A, Boulanger R, Davrieux F, Pinot M-N, Tardan E, Forestier-Chiron N, et al. Assessment of cocoa (Theobroma cacao L.) butter content and composition throughout fermentations. Food Research International. 2018;107: 675–682.

54. Afoakwa EO, Quao J, Takrama J, Budu AS, Saalia FK. Chemical composition and physical quality characteristics of Ghanaian cocoa beans as affected by pulp pre- conditioning and fermentation. J Food Sci Technol. 2013;50: 1097. doi:10.1007/S13197-011-0446-5

55. Hidayat T, Mulyawanti I. The usage of dried starter for re-fermentation of unfermented cocoa beans. IOP Conference Series: Earth and Environmental Science. IOP Publishing; 2019. p. 012061.

56. Bomdzele E, Molua EL. Assessment of the impact of climate and non-climatic parameters on cocoa production: a contextual analysis for Cameroon. Frontiers in Climate. 2023;5: 1069514. doi:10.3389/FCLIM.2023.1069514/BIBTEX

57. Chen X, Wu Y, Zhu H, Wang H, Lu H, Zhang C, et al. Turning over fermented grains elevating heap temperature and driving microbial community succession during the heap fermentation of sauce-flavor baijiu. LWT. 2022;172: 114173. doi:10.1016/J.LWT.2022.114173

58. JINAP S, DIMICK PS. Acidic Characteristics of Fermented and Dried Cocoa Beans from Different Countries of Origin. J Food Sci. 1990;55: 547–550. doi:10.1111/J.1365-2621.1990.TB06806.X

59. Jonfia-Essien WA, West G, Alderson PG, Tucker G. Phenolic content and antioxidant capacity of hybrid variety cocoa beans. Food Chem. 2008;108: 1155–1159.

60. Urbańska B, Kowalska J. Comparison of the Total Polyphenol Content and Antioxidant Activity of Chocolate Obtained from Roasted and Unroasted Cocoa Beans from Different Regions of the World. Antioxidants. 2019;8. doi:10.3390/ANTIOX8080283

61. Lund MN, Ray CA. Control of Maillard reactions in foods: Strategies and chemical mechanisms. J Agric Food Chem. 2017;65: 4537–4552.

62. Oracz J, Nebesny E. Effect of roasting parameters on the physicochemical characteristics of high-molecular-weight Maillard reaction products isolated from cocoa beans of different Theobroma cacao L. groups. European Food Research and Technology. 2019;245: 111–128.

63. Misnawi, Selamat J, Bakar J, Saari N. Oxidation of polyphenols in unfermented and partly fermented cocoa beans by cocoa polyphenol oxidase and tyrosinase. J Sci Food Agric. 2002;82: 559–566. doi:10.1002/JSFA.1075

64. Lin T, Chen L, Yang X, Fu C, Yuk H-G, Fan R, et al. Sensory Nutrition and Bitterness and Astringency of Polyphenols. Biomolecules 2024, Vol 14, Page 234. 2024;14: 234. doi:10.3390/BIOM14020234

65. Brillouet JM, Hue C. Fate of proanthocyanidins and anthocyanins along fermentation of cocoa seeds (Theobroma cacao L.). Journal of Applied Botany and Food Quality. 2017;90: 141–146. 10.5073/JABFQ.2017.090.017

66. Bauer D, de Abreu JP, Oliveira HSS, Goes-Neto A, Koblitz MGB, Teodoro AJ. Antioxidant activity and cytotoxicity effect of cocoa beans subjected to different processing conditions in human lung carcinoma cells. Oxid Med Cell Longev. 2016;2016.

67. Krähmer A, Engel A, Kadow D, Ali N, Umaharan P, Kroh LW, et al. Fast and neat– Determination of biochemical quality parameters in cocoa using near infrared spectroscopy. Food Chem. 2015;181: 152–159.

68. Mehrabani S, Arab A, Mohammadi H, Amani R. The effect of cocoa consumption on markers of oxidative stress: A systematic review and meta-analysis of interventional studies. Complement Ther Med. 2020;48: 102240. doi:10.1016/J.CTIM.2019.102240

69. Hu S, Kim B-Y, Baik M-Y. Physicochemical properties and antioxidant capacity of raw, roasted and puffed cacao beans. Food Chem. 2016;194: 1089–1094.

70. da Silva Oliveira C, Fonseca Maciel L, Spínola Miranda M, da Silva Bispo E. Phenolic compounds, flavonoids and antioxidant activity in different cocoa samples from organic and conventional cultivation. British Food Journal. 2011;113: 1094–1102.

71. Cuellar-Álvarez L, Cuellar-Álvarez N, Galeano-García P, Suárez-Salazar JC. Effect of fermentation time on phenolic content and antioxidant potential in Cupuassu (Theobroma grandiflorum (Willd. ex Spreng.) K. Schum.) beans. Acta Agron. 2017;66: 473–479.

72. Zzaman W, Bhat R, Yang TA. Effect of superheated steam roasting on the phenolic antioxidant properties of cocoa beans. J Food Process Preserv. 2014;38: 1932–1938.

73. Gu F, Tan L, Wu H, Fang Y, Xu F, Chu Z, et al. Comparison of Cocoa Beans from China, Indonesia and Papua New Guinea. Foods 2013, Vol 2, Pages 183–197. 2013;2: 183–197. doi:10.3390/FOODS2020183

74. ICCO. Who are the Fine and Flavour cocoa exporting countries? Abidjan, Cote d’Ivoire; 2020. Available: https://www.icco.org/fine-or-flavor-cocoa/

75. Romero JMV, Pabón YTM. Características sensoriales de granos y licor de cacao por un panel de jueces en entrenamiento. Revista SENNOVA: Revista del Sistema de Ciencia, Tecnología e Innovación. 2020;5: 27–42.

76. Martínez-Guerrero NC, Ligarreto-Moreno GA. Sensory analysis of cacao liquor (Theobroma cacao L.) in cultivars with different origins grown in the Colombian tropics. Revista Colombiana de Ciencias Hortícolas. 2023;17: e15876–e15876. doi:10.17584/RCCH.2023V17I2.15876

77. Portillo E, Labarca M, Grazziani L, Cros E, Assemat S, Davrieux F, et al. Formación del aroma del cacao Criollo (Theobroma cacao L.) en función del tratamiento poscosecha en Venezuela. Revista Científica UDO Agrícola. 2009;9: 458–468.

78. Erazo Solorzano CY, Disca V, Muñoz-Redondo JM, Tuárez García DA, Sánchez-Parra M, Carrilo Zenteno MD, et al. Effect of Drying Technique on the Volatile Content of Ecuadorian Bulk and Fine-Flavor Cocoa. Foods. 2023;12: 1065.

79. Nazario O, Ordoñez E, Mandujano Y, Arévalo J. Polifenoles totales, antocianinas, capacidad antioxidante de granos secos y análisis sensorial del licor de cacao (Theobroma cacao L.) criollo y siete clones. RevIA. 2018;3.

80. Rodriguez-Campos J, Escalona-Buendía HB, Orozco-Avila I, Lugo-Cervantes E, Jaramillo-Flores ME. Dynamics of volatile and non-volatile compounds in cocoa (Theobroma cacao L.) during fermentation and drying processes using principal components analysis. Food Research International. 2011;44: 250–258.

81. Cedeño P. Determinación de perfiles organolépticos en ocho grupos de cacao mediante la degustación de licor de cacao y chocolates oscuros elaborados artesanalmente. Calceta, Ecuador Carrera de Agroindustria Escuela Superior Politécnica Agropecuaria de Manabí. 2010.

82. Machado Cuellar L, Ordoñez Espinosa CM, Angel Sánchez YK, Guaca Cruz L, Suárez Salazar JC. Organoleptic quality assessment of Theobroma cacao L. in cocoa farms in northern Huila, Colombia. Acta Agron. 2018;67: 46–52.

83. de Andrade AB, da Cruz ML, de Souza Oliveira FA, Soares SE, Druzian JI, de Santana LRR, et al. Influence of under-fermented cocoa mass in chocolate production: Sensory acceptance and volatile profile characterization during the processing. Lwt. 2021;149: 112048.

84. Tee Y, Bariah K, Hisyam Zainudin B, Samuel Yap K, Ong N. Impacts of cocoa pod maturity at harvest and bean fermentation period on the production of chocolate with potential health benefits. J Sci Food Agric. 2022;102: 1576–1585.

85. Rodríguez Silva LG, Quintana Fuentes LF, Coronado Silva RA, García Jerez A, Baez Daza EY, Agudelo-Castañeda GA. Caracterización física y sensorial de 24 genotipos especiales de cacao Theobroma cacao. Revista UDCA Actualidad & Divulgación Científica. 2023;26: 1–11.

86. Santander Muñoz M, Rodríguez Cortina J, Vaillant FE, Escobar Parra S. An overview of the physical and biochemical transformation of cocoa seeds to beans and to chocolate: Flavor formation. Crit Rev Food Sci Nutr. 2020;60: 1593–1613.

